# Single-cell transcriptome unravels spermatogonial stem cells and dynamic heterogeneity of spermatogenesis in seasonal breeding teleost

**DOI:** 10.1101/2024.06.08.598045

**Authors:** Yang Yang, Yinan Zhou, Gary Wessel, Weihua Hu, Dongdong Xu

## Abstract

Seasonal spermatogenesis in fish is driven by spermatogonial stem cells (SSCs), which undergo a complex cellular process to differentiate into mature sperm. In this study, we characterized spermatogenesis in the large yellow croaker (*Larimichthys crocea*), a marine fish of significant commercial value, based on a high-resolution single-cell RNA-seq atlas of testicular cells from three distinct developmental stages- juvenile, adult differentiating and regressed testes. We detailed continuous developmental trajectory of spermatogenic cells, from spermatogonia to spermatids, elucidating the molecular events involved in spermatogenesis. We uncovered dynamic heterogeneity in cellular compositions throughout the annual reproductive cycle, accompanied by strong molecular signatures within specific testicular cells. Notably, we identified a distinct population of SSCs and observed a critical metabolic transition from glycolysis to oxidative phosphorylation, enhancing our understanding of the biochemical and molecular characteristics of SSCs. Additionally, we elucidated the interactions between somatic cells and spermatogonia, illuminating the mechanisms that regulate SSCs development. Overall, this work enhances our understanding of spermatogenesis in seasonal breeding teleost and provides essential insights for the further conservation and culture of SSCs.

**Summary statement:** Our study reveals new insights into the development of spermatogonial stem cells (SSCs), potentially impacting further conservation and culture of SSCs in teleost.

## Introduction

Spermatogenesis is a highly complex process, during which spermatogonia differentiate into mature sperm through sequential stages of mitosis and meiosis (Schulz et al., 2010; Yoshida, 2016). Spermatogonial stem cells (SSCs), the only germline stem cells, are a rare subset of spermatogonia and capable of dual abilities of both self-renewal and differentiation (Kubota and Brinster, 2018; Lacerda et al., 2014). As the foundation units of continual spermatogenesis, the normal development of SSCs is critical not only for successful fertility but also ensuring the continuation of species in vertebrates. Traditional approaches to studying spermatogenesis have predominantly relied on histological observation and several specific gene analyses, which have provided limited insights into the developmental transitions of spermatogonia in vertebrates (Oatley and Brinster, 2006; von Kopylow and Spiess, 2017). The molecular mechanisms governing SSCs development and their regulatory remain poorly understood, particularly in non-mammalian vertebrates such as fish.

Fish, representing the most diverse group of vertebrates, display unique and varied patterns of spermatogenesis that are significantly influenced by environmental factors. In most teleost species, spermatogenesis is both seasonal and cyclic, characterized by profound morphological and cellular compositional changes in the germinal epithelium throughout the reproductive cycle (Chaves-Pozo et al., 2005; Hernández-Franyutti and Uribe, 2012). Initially, in juvenile males, the testes remain quiescent, characterized by an abundance of undifferentiated spermatogonia, including SSCs. The stemness of these undifferentiated spermatogonia has been confirmed through germ cell transplantation method in various fish species, such as in rainbow trout (*Oncorhynchus mykiss*) (Sato et al., 2017), medaka (*Oryzias latipes*) (Seki et al., 2017), turbot (*Scophthalmus maximus*) (Zhou et al., 2021) and blue drum (*Nibea mitsukurii*) (Xu et al., 2019). From the onset of puberty, these undifferentiated spermatogonia begin continuously entry into spermatogenesis, cycling through developmental stages in adult testes (Sato et al., 2017; Yang et al., 2018; Yu et al., 2024). At postspawning, the testes enter into a regressed process, during which diploid spermatogonia predominantly remain as the germ cells capable of initiating the next cycle of spermatogenesis (Sato et al., 2017; Yu et al., 2024). This rigorous spatiotemporal orchestration of spermatogenesis provides a critical framework for analyzing the characteristics and regulatory mechanisms of SSCs development in fish. Understanding the mechanisms of fish SSCs development not only provides insights into the dynamics of spermatogenesis in seasonal breeding fish, but also establishes a foundation for the effective utilization and conservation of SSCs in teleost.

The advent of single-cell RNA sequencing (scRNA-seq) has opened new avenues for unraveling the complex gene regulatory networks underlying spermatogenesis. Recent studies in mammals have created comprehensive testicular transcriptional cell atlases and identified molecular markers for different testicular cell types (Dong et al., 2023; Guo et al., 2018; Hermann et al., 2018; Wei, et al., 2021). Although similar studies in several teleosts, such as zebrafish (*Danio rerio*) (Qian et al., 2022), black rockfish (*Sebastes schlegelii*) (Wang et al., 2023) and Chinese tongue sole (*Cynoglossus semilaevis*) (Wang et al., 2022), have begun to sketch the testicular cell landscape, our understanding of fish SSCs and testicular development during the annual reproductive cycle remains incomplete.

The large yellow croaker (*Larimichthys crocea*), a member of family Sciaenidae, is a seasonally breeding species and one of the most commercially valuable marine fishes in the East Asia (Wu et al., 2014; Yan et al., 2022). Historically, it was among the top three commercial marine species in China. However, by the late 1980s, the species had become “threatened” due to overfishing and environmental pollution (Chen et al., 2018; Yu et al., 2023). As the most extensively cultured marine species in China, the aquaculture of large yellow croaker has spread rapidly along the southeast coast, driven by its high nutritional value (Chen et al., 2018). This growth significantly contributes to the economy, offering a vital source of protein and generating substantial income for millions of people. Understanding the molecular basis of spermatogenesis, particularly the development of SSCs in this species, is of great importance not only for enhancing its aquaculture sustainability through SSCs biotechnology, but also for gaining insights into the reproductive strategies of other fish species. In this study, we performed scRNA-seq on testicular tissues from large yellow croaker at distinct developmental stages, including juvenile testes and adult differentiating and regressed testes. Our findings offer a comprehensive testicular transcriptional cell atlas, suggesting dynamic stage-specific heterogeneity for spermatogenesis in seasonal breeding teleost. We further delineate the characteristics of a novel SSCs state and their regulatory mechanisms, uncovering the first molecular signatures for SSCs in teleost.

## Results

### 1. ScRNA-seq identifies major germ and somatic cell types of testes within large yellow croaker

The scRNA-seq of testes from large yellow croaker were established under the 10× Genomics using the droplet-based methodology (Fig.1A). We have detailly analyzed the testicular development during the annual reproductive cycle using histological observation. According to the main and typical germ cells, we classified the testes into six developmental stages (from I to VI stages, Fig.1B), as follows: (I) spermatogonia proliferation stage. This stage occurs only once in the life of a fish. Some spermatogonia (SPG) with a prominent and clear nucleus were dispersed in the testes (Fig.1Ba). (II) early spermatogenesis stage. A large number of SPG were observed, and some scattered primary spermatocyte (PSPC) and secondary spermatocyte (SSPC) cysts were also observed in the testes (Fig.1Bb). (III) late spermatogenesis stage. Many cysts containing differentiating germ cells were prominent in the testes, a large number of PSPC and SSPC and some spermatozoa (E) were dispersed in the lumen of the testicular lobules (Fig.1Bc). (IV) early spermiation stage. A large number of E were released into the lumen of the testicular lobules, and some spermatogenic cysts could still be observed along the walls of the spermatogenic lobules (Fig.1Bd). (V) late spermiation stage. The testis was filled with a large number of E (Fig.1Be). (VI) regression stage. The residual E and some SPG were observed in the testes (Fig.1Bf). In addition, we further performed the immunofluorescence of Vasa (a well-known germ cell marker) in the testes at six stages (Fig.1C). We found that Vasa prominently expressed in SPG. With spermatogenesis, the signals of Vasa became weak, and no signal was detected in secondary spermatocyte (SSPC) and E. Regarding Vasa abundance in testes, significant signals were observed in the testes at stages I, II and VI.

**Figure 1.**
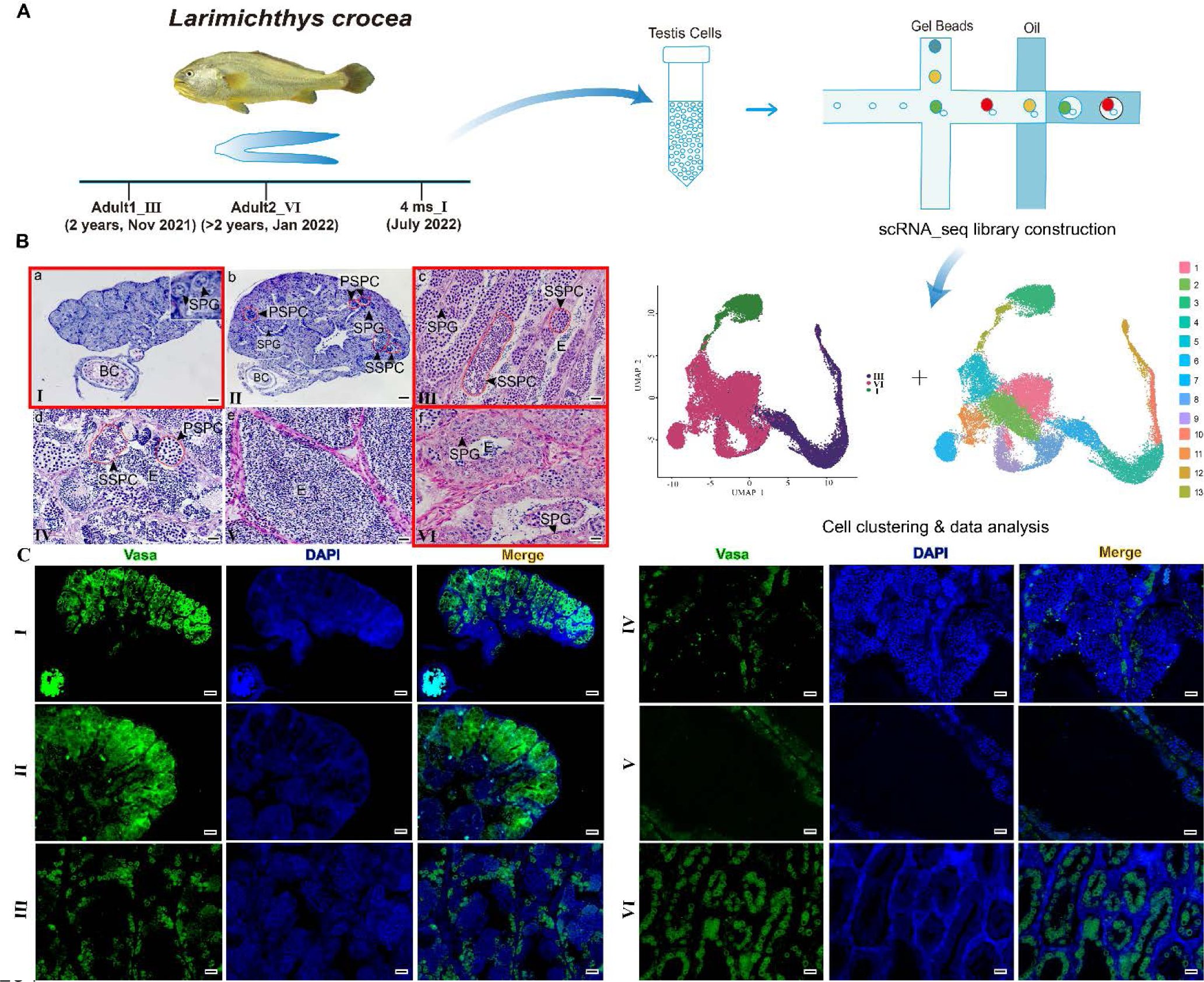
Single-cell transcriptome profiling of large yellow croaker testes at distinct developmental stages. (A) Schematic showing single-cell RNA-seq of large yellow croaker testicular cells at three developmental stages. (B) H&E staining of testes at different developmental stages from I to VI. Scale bars represent 20 μm. (C) Immunofluorescence of Vasa (green) in the testes at different developmental stages from I to VI. Nuclei were visualized using DAPI (blue). Scale bars represent 20 μm. BC, blood capillary; SPG, spermatogonia; PSPC, primary spermatocyte; SSPC, secondary spermatocyte; E, spermatozoa.

In order to identify all the types of germ cells, especially the SPG in testes, we created 3 scRNA-seq datasets from the testes at stages I, III and VI. Among them, testicular cells from ∼50 fishes (4-months old) were pooled for constructing the dataset of testes at stage I. The datasets of testes at stage III and VI were accomplished from two fish obtained in the Autumn (November) and Winter (January) respectively. After standard quality control (QC) dataset filters, we obtained a total of ∼3,7370 cells for transcriptional spectra construction. To minimize the effects from batch-related processing of samples, we integrated all 3 scRNA-seq datasets using the mutual nearest neighbor (MNN) methodology. Each cell had an average number of 5552 unique molecular identifiers (UMI) and 2606 genes detected. We next performed dimensionality reduction using t-distributed stochastic neighbor embedding (t-SNE) and cell clustering using shared nearest neighbor (SNN) methodology. In final, the transcriptional spectra were assigned to 13 cell clusters on unsupervised clustering and visualized by uniform manifold approximation and projection (UMAP) analysis (Fig.1A).

Based on known cell-type marker genes and differentially expressed genes (DEGs), 13 testicular cell clusters were identified and clarified into 11 major cell types, including 8 germ cell clusters and 3 somatic cell types (Figure 2A and B). Germ cell types were identified through markers as follows: undifferentiated spermatogonia (Undiff SPG, marked by *pou5f3* and *cd9*), differentiated spermatogonia (Diff SPG, marked by *dnd1* and *daz1*), preleptotene spermatocyte (Pre-Lep, marked by *cdca7*, *mcm6* and *mcm3*), pachytene spermatocyte (P-SPC, marked by *ccnb3* and *smc4*), meiotic spermatocyte (M-SPC, marked by *dmc1* and *meiob*), initial spermatids (E1, marked by *nme5*), intermediate spermatids (E2, marked by *dnah3*) and final spermatids (E3, marked by *tekt1*) (Figure 2C and D). In addition, we also identified 3 somatic cell types as Sertoli cell (marked by *gsdf* and *inha*), Leydig cell (marked by *igf2* and *insl3*) and NKT cell (marked by *cxcr4* and *impa1*) (Figure 2C and D). Vasa was expressed by all germ cell types (Figure 2C). For validation, we performed in *situ* hybridization (ISH) analysis on some cell marker genes, including *pou5f3, dnd1, dmc1, akap14* and *gsdf* (Figure 2E). We found that *pou5f3* were expressed in Undiff SPG at the periphery of the testes, while *dnd1* were transcribed in Diff SPG. Notably, *dmc1* and *akap14* were respectively expressed in M-SPC and sperm. The Sertoli cell marker gene *gsdf* was obviously expressed in somatic cells around spermatogonia. These genes provide new resources as candidate marker genes for each cell type. Altogether, our atlas captures the majority of testicular cell types and allows us to directly compare gene expression patterns among the testis cell types.

**Figure 2.**
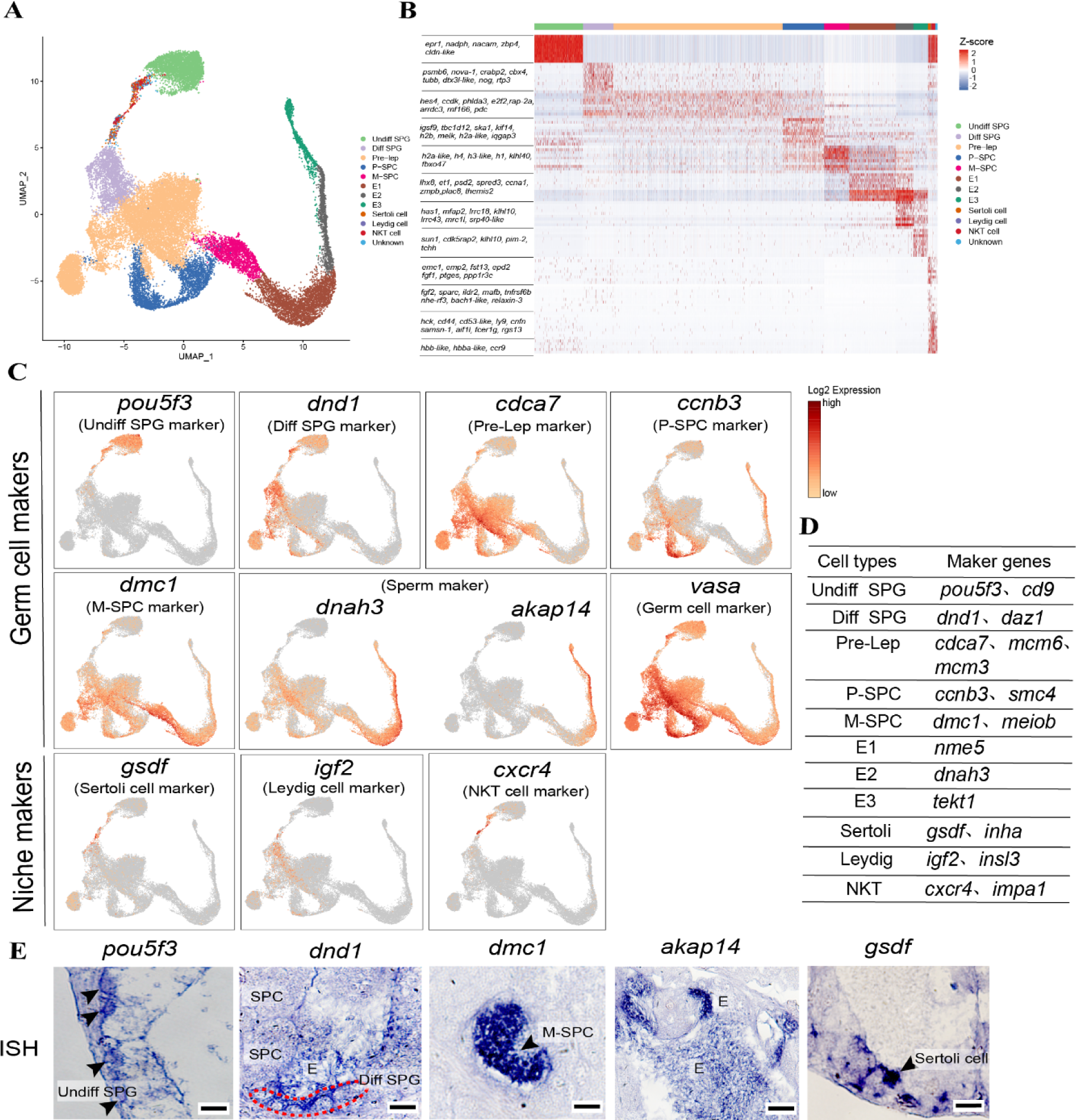
Single-cell RNA-seq establishes germ and somatic cell types in large yellow croaker testes. (A) A UMAP plot showing the cell types from large yellow croaker testes. Each dot represents a single cell and is colored according to its cluster identity. (B) Heatmap showing the top 20 DEGs for 11 cell types. (C) Expression patterns of selected marker genes on the UMAP plot. Red (or orange) represents a high (or low) expression level. (D) A table of maker genes for various cell types. (E) *In situ* hybridization of selected marker genes in large yellow croaker testes. Positive signals are indicated by arrows and colored with blue. Scale bars represent 30 μm. Undiff SPG, undifferentiated spermatogonia; Diff SPG, differentiated spermatogonia; Pre-Lep: preleptotene spermatocyte; P-SPC, pachytene spermatocyte; M- SPC, meiotic spermatocyte; E1, initial spermatids; E2, intermediate spermatids; E3, final spermatids.

### 2. Developmental trajectory and dynamic gene expression patterns during spermatogenesis in large yellow croaker

To further assess the developmental trajectory, we performed pseudotime analysis of all germ cell types using Monocle2. We found the directionality along pseudotime precisely correlated with the trajectory of spermatogenesis determined by our initial germ cell cluster analysis (Figure 3A). Using the trajectory from SPG through SPC to E, we analyzed the expression patterns of genes during spermatogenesis, revealing four distinct gene cohorts (Supplementary Figure 1A). We found a large number of genes were highly expressed during spermiogenesis (Supplementary Figure 1A). We further analyzed the expression of marker genes in Undiff SPG and Diff SPG. Noticeable transitions in gene expression as visualized on the heatmap, corresponded with key cell-state-transitions from Undiff SPG to Diff SPG (Figure 3B). For instance, we identified genes involved in Undiff SPG (e.g., *pou5f3, cd9, klhdc1, arhgef19, lsm14b, snx10a, rgs14* and *foxgla*) and genes involved in Diff SPG (e.g., *dnd1, id3, id1, irf1, smad9* and *mf19b*). We found that Undiff SPG expressed more genes involved in the maintenance of stemness and began to express genes related to spermatogonia differentiation in Diff SPG. In addition, we also analyzed cell-cycle related genes expression in diverse germ cell types. Diff SPG and Pre-Lep expressed a number of G1/S genes (e.g., *mcm4, mcm5, mcm6, msh6, hells, smarcad1* and etc.), whereas P-SPC highly expressed G2/M genes (e.g., *cdk1, ccnb3, cdca3* and etc.) and M-SPC expressed M staged genes (e.g., *ccdc36, meiob, sycp3, dmc1, sycp2* and etc.) (Figure 3C).

**Figure 3.**
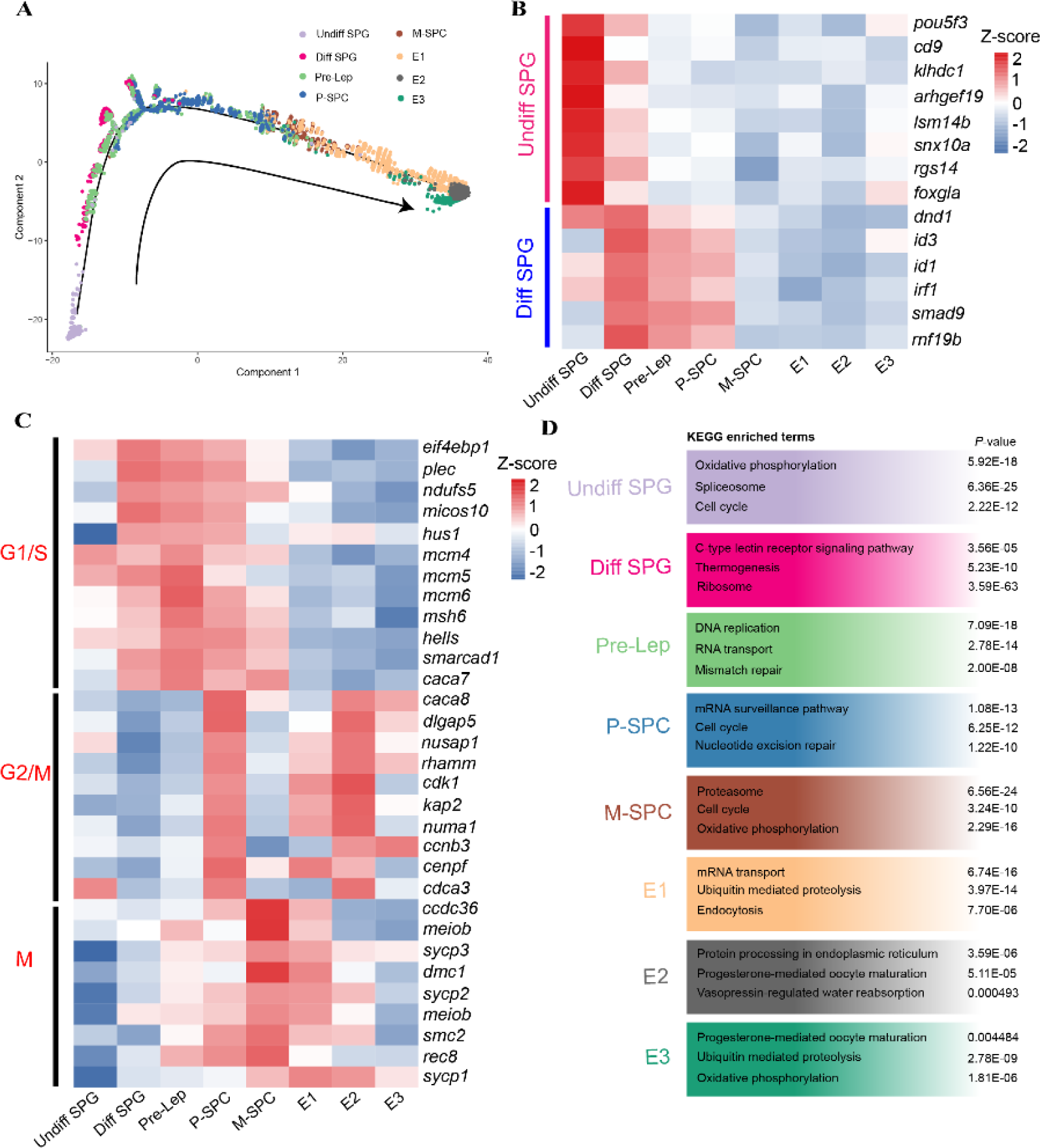
Analysis of germ cell development and spermatogenesis trajectory in the large yellow croaker. (A) Pseudotime analysis of 8 germ cell types. Different germ cells were annotated with distinct colors. (B) Heatmap of DEGs of Undiff SPG and Diff SPG in 8 germ cell types. (C) Heatmap of cell cycle- specific genes in 8 germ cell types. (D) Functional enrichment analysis of DEGs in 8 germ cell types. Undiff SPG, undifferentiated spermatogonia; Diff SPG, differentiated spermatogonia; Pre-Lep: preleptotene spermatocyte; P-SPC, pachytene spermatocyte; M-SPC, meiotic spermatocyte; E1, initial spermatids; E2, intermediate spermatids; E3, final spermatids.

We then performed functional enrichment analysis of DEGs in each germ cell type (Figure 3D). According to the KEGG analysis, DEGs in Undiff SPG were mainly enriched in “oxidative phosphorylation”, “cell cycle” and “spliceosome”; DEGs in Diff SPG were mainly enriched in “c-type lectin receptor signaling pathway”, “thermogenesis” and “ribosome”; DEGs in Pre-Lep were mainly enriched in “DNA replication”, “RNA transport” and “mismatch repair”; DEGs in P-SPC were mainly enriched in “cell cycle”, “oxidative phosphorylation” and “proteasome”; DEGs in E1 were mainly enriched in “mRNA transport”, “ubiquitin mediated proteolysis” and “endocytosis”; DEGs in E2 were mainly enriched in “protein processing in endoplasmic reticulum”, “progesterone-mediated oocyte maturation” and “vasopressin-regulated water reabsorption”; DEGs in E3 were mainly enriched in “progesterone-mediated oocyte maturation”, “ubiquitin mediated proteolysis” and “oxidative phosphorylation” (Figure 3D).

To further obtain a detail view of spermiogenesis, we analyzed expression of DEGs in three sperm clusters (Supplementary Figure 1B and C). There were obvious differences in gene expression among three clusters. We found more genes that play crucial roles in cilia formation were highly and specifically expressed in E2, such as coiled-coil domain containing protein (*ccdc138, ccdc178* and *ccdc39*), cilia and flagella associated protein (*cfap43, cfap47, cfap69, cfap70*) and cilia ODA dynein axonemal heavy chain (DNAH) protein (*dnah1, dnah3, dnah6, dnah7, dnah8, dnah10*). Cell-cycle related genes were specifically expressed in E1, including synaptonemal complex protein (*sycp1, sycp2* and *sycp3*) and cyclin N-termial domain-containing 1 (*cntd1*). For E3, the genes essential for sperm flagellar motility were specifically expressed, such as tektin1 (*tekt1*), sperm-associated antigen 6 (*spag6*) and a kinase anchor protein (*akap14*). Furthermore, we analyzed the dynamic expression of some genes (*cystm1, tekt1, tspan7, hydin, spag1* and *inf2*) during spermiogenesis and found their expression in different clusters (Supplementary Figure 1B).

### 3. Different proportions of cell types in testes at different developmental stages

We further examined the distribution of cell types and quantified their relative proportions in the testes at different stages (Figure 4A and B). Notably, most Undiff SPG were detected in the testis at stage I. Somatic cells (Sertoli cells, Leydig cells and NKT cells) were also mainly detected in the testis at stage I. Nevertheless, Diff SPG, spermatocytes and sperm were detected in the testes at stages III and VI. In comparison to Undiff SPG, Diff SPG and early spermatocytes (Pre-lep and P- SPC) were observed in the testis at stage VI. Cell types of spermatocytes and sperm were mainly detected in the testis at stage III. In addition, pseudotime analysis showed the developmental trajectory was from stage I through VI to III (Figure 4C). Those results indicated that testicular cell composition was heterogeneously impacted by developmental stages, consistent with previous histological examination.

**Figure 4.**
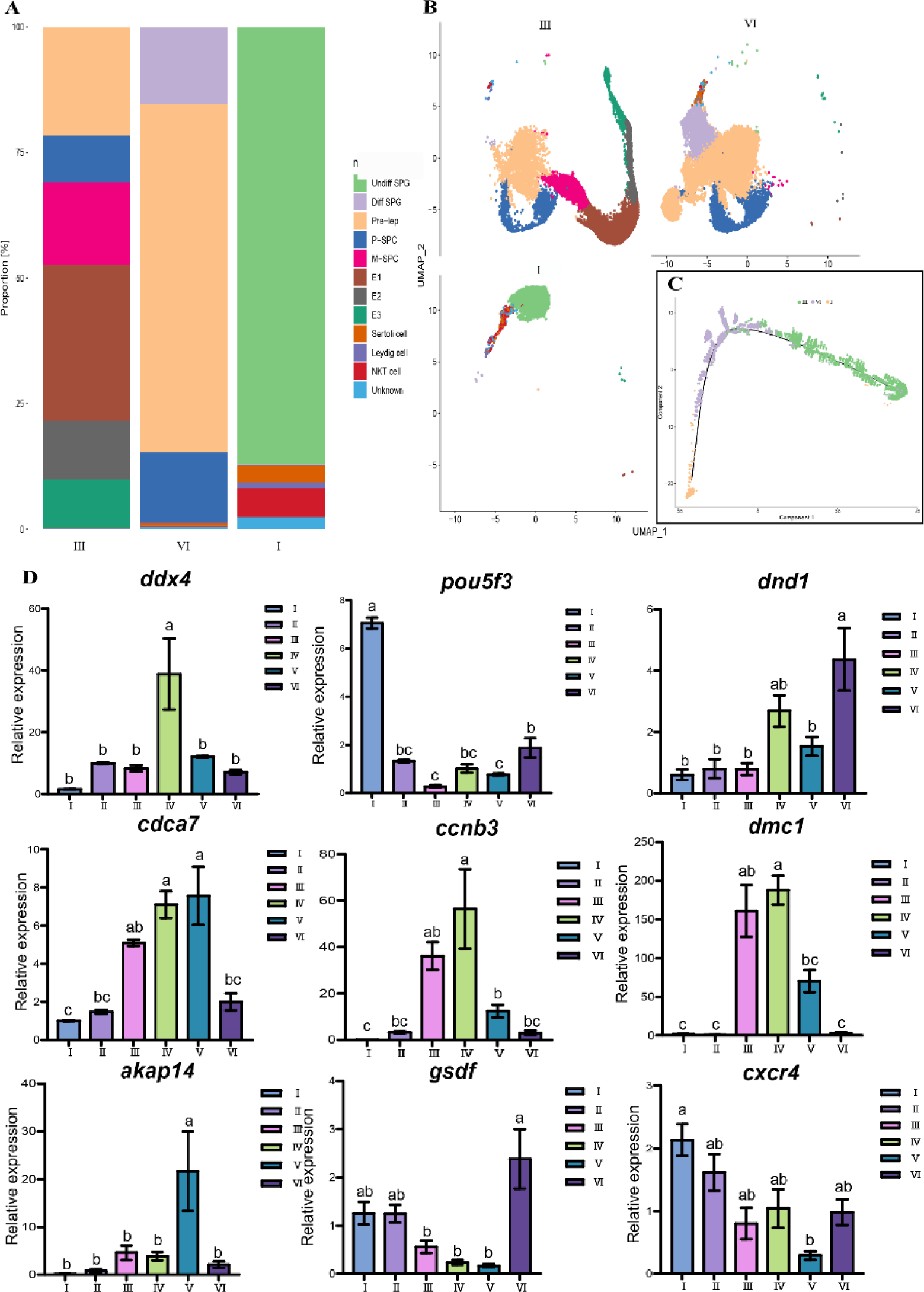
Analysis of distribution of cell types and selected marker genes expression. (A) Histogram of testicular cell types at distinct developmental stages. (B) Distribution of testicular cells at distinct developmental stages on the UMAP plot. (C) Pseudotime analysis of testicular cells at distinct developmental stages were colored with different colors. (D) Expression of selected marker genes at six developmental stages by RT-qPCR. Different testicular developmental stages were colored with different colors. Undiff SPG, undifferentiated spermatogonia; Diff SPG, differentiated spermatogonia; Pre-Lep: preleptotene spermatocyte; P-SPC, pachytene spermatocyte; M-SPC, meiotic spermatocyte; E1, initial spermatids; E2, intermediate spermatids; E3, final spermatids.

We then performed RT-qPCR to verify the results of scRNA-seq using various germ and somatic cells-specific marker genes, namely *vasa, pou5f3, dnd1, cdca7, ccnb3, dmc1, akap14, gsdf* and *cxcr4* (Figure 4D). The Undiff SPG maker *pou5f3* demonstrated a high level of expression in the testes at stage I, diminished expression from stage II to V, and significantly increased transcripts at stage VI. Diff SPG marker *dnd1* highly expressed in testes at stage VI. The SPC markers *cdca7, ccnb3* and *dmc1* were mainly expressed in testes at stages III and IV, whereas the sperm marker *akap14* was abundantly detected in testis at stage V, reflecting active spermatogenesis in testes from stages III to V. *Vasa*, a marker gene of SPG and PSPC, was obviously expressed in testes at IV stage. The NKT marker *cxcr4* and Sertoli cell marker *gsdf* was abundantly expressed in testes at stages IV, I and II. Together these results corroborated the scRNA-seq results and illustrated that the expression levels of various germ and somatic cells-specific marker genes varied significantly depending on the reproductive stages, reflecting the dynamic heterogeneity of spermatogenesis.

### 4. Identification of spermatogonial stem cells (SSCs) in large yellow croaker

To further resolve the spermatogonial ‘states’ in order to understand the development progression of spermatogonia, we further profiled testicular cells from the juvenile (4-ms old) testes. After QC filtering, we obtained ∼4742 single cells and assigned cell clusters based on known markers (Supplementary Figure 2A and Figure 5B). This analysis identified 3 somatic cells (i.g., Sertoli/Leydig, Erythrocyte and Leukocyte), and 5 germ cell types (Figure 5A and B). Three germ cell clusters showed high similarity to the Undiff SPG previously described, hereafter we termed them as Undiff SPG state 0, state1 and state2 (Figure 5A and B). Notably, we analyzed and observed high expression of markers of Undiff SPG *(arhgef19, lsm14b, klhdc1, cd9* and *pou5f3*) in a small portion of cells in Undiff SPG state 0 (Supplementary Figure 2B). Therefore, we speculated that those cells (< 30 cells) may represent the earliest/naïve SSCs (Figure 5C). Further, we performed RNA velocity analysis, a computational approach that utilizes nascent transcription to infer developmental trajectories. Within each cell, the amplitude and direction of the vector reflects a transcriptional trajectory. Notably, we found SSCs displayed long velocity vectors turning into a state that was likely to be stem cells (Figure 5D). We also observed a sub-group of Undiff SPG contained long vectors, indicating an apparent progression towards SPC whereas most Undiff SPG lacked long vectors. We then performed pseudotime analysis and observed the predicted SSCs located in the start of trajectory of Undiff SPG (Supplementary Figure 2C and Figure 5D). This pattern suggests that the classified SSC is potentially a spermatogonial stem cell.

**Figure 5.**
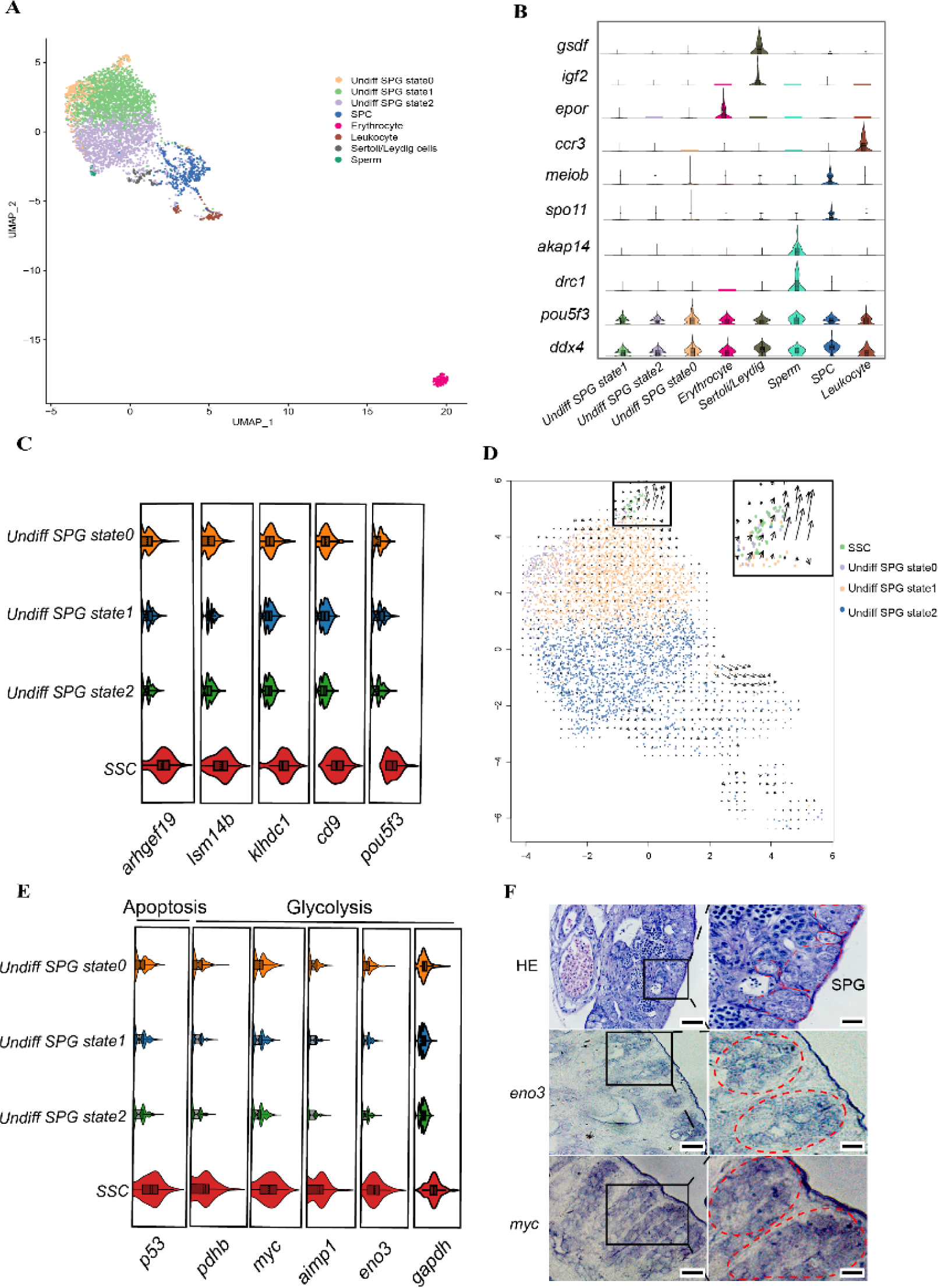
Single-cell transcriptome profiling and analysis from infant testes (4-ms old) of large yellow croaker. (A) A UMAP plot showing 8 cell types from infant testes. (B)Violin plots of the normalized expression of marker genes for the 8 cell types. (C)Violin plots of Undiff SPG marker genes in SSC and Undiff SPG states 0-2. (D)Visualization of the RNA velocity analysis on UMAP plot. (D)Violin plots of apoptosis gene (*p53*) and glycolysis genes (*pdhb*, *myc*, *aimp1*, *eno3* and *gapdh*) in SSC and Undiff SPG stage 0-2. (E) *In situ* hybridization of glycolysis genes (*eno3* and *myc*) in large yellow croaker testes. The right pictures are magnified of the black wireframes of left pictures. Red dashed box represents the SPG regions. Undiff SPG, undifferentiated spermatogonia; Diff SPG, differentiated spermatogonia; SPC, spermatocyte.

We then performed functional enrichment analysis of markers of SSC and undiff SPG (state 0- 2). Genes in SSC were enriched in “cellular response to DNA damage stimulus” and “mitochondrial matrix” (i.e., *sirt4, atad5b, caspase3, casp9* and etc); genes in Undiff SPG state 0 were enriched in “respiratory chain” and “respiratory chain complex” (i.e., *nd1, nd2, nd3, cox3* and etc); genes in Undiff SPG state 1 and 2 were enriched in “histone deacetylase binding”, “cyclin-dependent protein serine/threonine kinase regulator activity” and “DNA replication initiation” (i.e., *cdc20, cdc45, mcm7, cyclin-B2* and etc) (Supplementary Figure 2E). This analysis also identified metabolic differences between SSC and Undiff SPG. We were excited to find that SSC had high gene expressions of glycolysis (i.g., *myc, aimpa, eno3* and *gaphd*) and apoptosis (*p53*), compared with Undiff SPG (Figure 5E). On the other hand, we also observed the Undiff SPG state 0 displaying higher expression of those genes than the Undiff SPG state 1 and 2, indicating the Undiff SPG state 0 presents a state between SSC and Undiff SPG. For validation, we performed *in situ* hybridization (ISH) on *eno3* and *myc* gene. In testes, SPGs were mainly located in the edge and displayed high expression of *eno3* and *myc* (Figure 5F). We then analyzed the DEGs and identified a number of potential marker genes of SSC (i.e., *ly75-like, bzwz, nthl1* and *cd200*) (Supplementary Figure 2F). To further verify the potential marker genes of SSC, we used ISH to validate the expression patterns of *ly75-like* and *ox2*. We found that *ly75-like* and *ox2* were highly expressed in Undiff SPG located in the edge of testes, while they were also slightly expressed in Diff SPG (Figure 6A).

**Figure 6.**
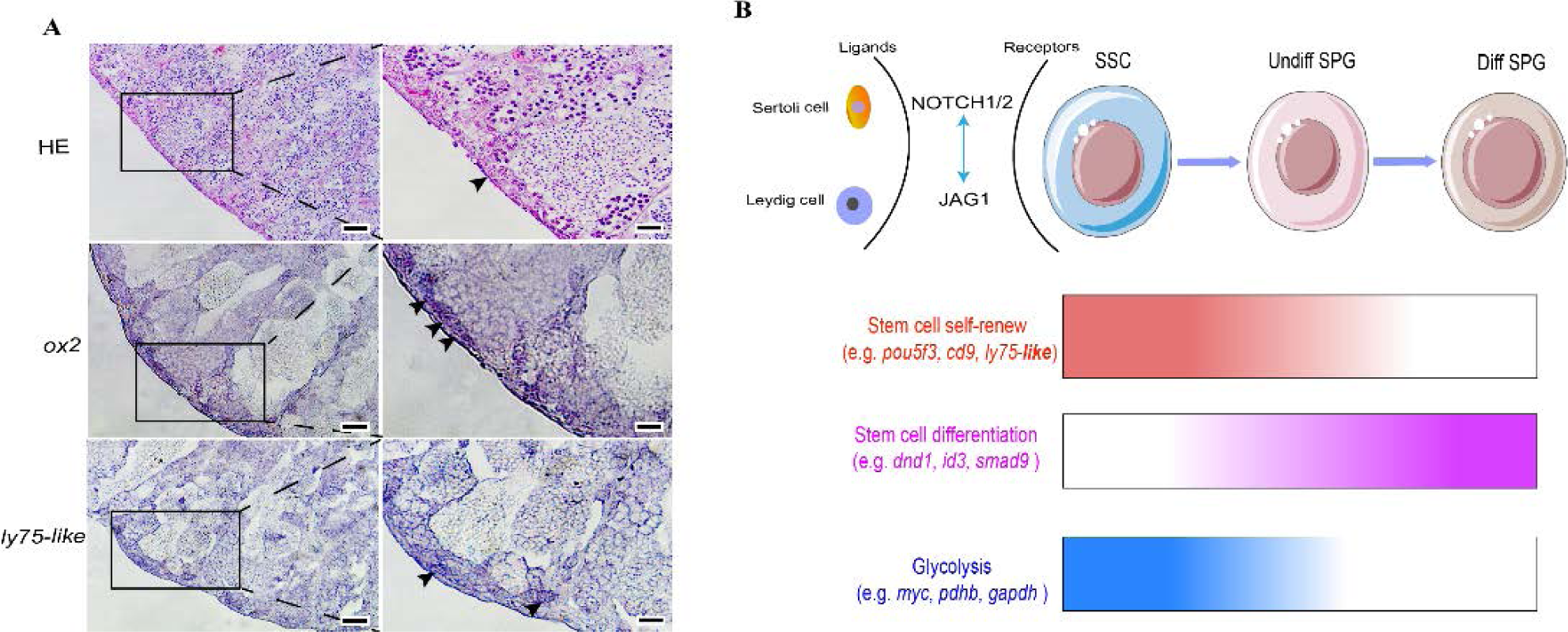
In situ hybridization of SSC marker genes and schematic illustrating the spermatogonia development and the interaction with somatic cells of large yellow croaker. (A) *In situ* hybridization of SSC marker genes (*ox2* and *ly75-like*) in large yellow croaker testes. Positive signals are indicated by arrows and colored with blue. The right pictures are magnified of the black wireframes of left pictures. Scale bars represent 50 μm. (B) Schematic illustrating the spermatogonia development and the interaction with somatic cells. the spermatogonia development from SSC to Diff SPG is regulated by Sertoli and Leydig cells by NOTCH1/2-JAG1 system. The marker genes for spermatogonia subsets are indicated.

### 5. Somatic cells and the crosstalk between somatic cells and germ cells from the juvenile testes

Sertoli cells and Leydig cells play supporting roles in spermatogenesis. Interestingly, we found the specific marker genes (*gsdf* for Sertoli cell and *igf2* for Leydig cell) highly expressed in the same cell cluster from the juvenile testes (Figure 5A and B). We named the cell cluster as Sertoli/Leydig cells. To further distinguish Sertoli cells and Leydig cells, we performed a focused analysis of Sertoli/Leydig cells using more marker genes, including *gsdf* and *fgf2* for Sertoli cells, *igf2* and *bmp4* for Leydig cells (Supplementary Figure 3A). We found that some cells present high expression levels of both the Sertoli and Leydig cell marker genes. The results suggested that Sertoli cells and Leydig cells may be differentiated from the same precursor cell type (Supplementary Figure 3A).

Subsequently, we analyzed the communication among various testicular cells using Cellphone B from the juvenile testicular cells. Among all cell types, Sertoli/Leydig cells exhibited robust interaction with four subsets of Undiff SPG (SSC and Undiff SPG state 0-2) (Supplementary Figure 3B). Notably, NOTCH1/2 tightly bound to JAG1 between Sertoli/Leydig cells and Undiff SPG, suggesting that Sertoli/Leydig cells play pivotal roles in spermatogonia maintenance and development. In addition, we found the potential for robust communications in Sertoli/Leydig cells through four ligand-receptor pairs, including PTPRK-BMP7, PLXNB2-SEMA4C, NOTCH1-JAG1 and NOTCH2-JAG1 (Supplementary Figure 3B).

## Discussion

Fish spermatogenesis is a cyclic and seasonal process involving the complex differentiation of SSCs into mature sperm. Throughout the annual reproductive cycle, the germinal epithelium undergoes significant changes, displaying distinct morphological characteristics and cellular compositions at different development stages. However, the cellular and molecular basis underlying the stage-specific heterogeneity of fish testicular cells, particularly the development of SSCs, is not understood. In this study, we performed scRNA-seq profiling of testicular tissues from large yellow croaker at three distinct developmental stages: juvenile, adult differentiating, and regressed testes. Significant changes in cellular compositions were observed in different developmental stages, accompanied by strong molecular signatures within specific testicular cells. Notably, we identified a distinct population of SSCs, and observed a significant metabolic shift from glycolysis to oxidative phosphorylation, along with niche-guided signaling pathways regulated by somatic cells that influence SSCs development. Our findings provide the first discovery of key molecular signatures essential for fish SSCs development, offering crucial insights for the further conservation and culture of SSCs in teleost.

### 1. Comprehensive cellular and molecular map of testicular development

Using scRNA-seq, we delineated 11 major cell types, including 8 germ cell clusters and 3 somatic cell types in large yellow croaker. The major cell types and typical marker genes in testes were similar with other fishes (Qian et al., 2022; Wang et al., 2022; Wang et al., 2023; Wu et al., 2021). In an unsupervised pseudotime analysis, the development directionality of germline cells was from SPG through SPC to E followed a continuous trajectory. The development trajectory is consistent with the process of spermatogenesis from teleost to mammals (Schulz et al., 2010). These results indicate that the process of spermatogenesis is highly conserved in vertebrates, wherein spermatogonia sequentially differentiate into primary spermatocytes, secondary spermatocytes, spermatids and ultimately mature into spermatozoa.

Furthermore, we identified novel markers for SPG (e.g., *pou5f3, cd9, arhgef19, lsm14b*, and *klhdc1* for Undiff SPG and *dnd1, smad9, mf19b* and *id1* for Diff SPG), some of which were also commonly highly expressed in spermatogonia of human and mice (Persio and Neuhaus, 2023; Dong et al., 2023; Itman and Loveland, 2013; Kanatsu-Shinohara et al., 2004; Niimi et al., 2019; Yokota, 2001). We also found that Undiff SPG expressed more genes involved in the maintenance of stemness and began to express genes related to differentiation in Diff SPG. Our data also provided a molecular map of germline cells for the cell cycle, demonstrating that germ cells were stayed in different rephrases for meiosis and mitosis. The results are consistent with the process of spermatogenesis using histological observation. In addition, we further analyzed three sperm clusters, and found a large number of spermiogenesis related genes, such *ccdcs, cfaps* and *dnahs*. Those genes also play essential roles in spermiogenesis in mammals (Levkova et al., 2022; Priyanka and Yenugu, 2021; Tang et al., 2017), indicating that the essential genes for spermiogenesis are conserved in vertebrates.

### 2. Stage-specific heterogeneity of spermatogenic cell during spermatogenesis

To investigate the heterogeneity in spermatogenesis, we examined the distribution and quantified the relative proportions of different cell types in the testes at various developmental stages. Notably, Undiff SPG and somatic cells (Sertoli cells, Leydig cells and NKT cells) were mainly detected in the testes at stage I, while Diff SPG, spermatocytes and sperm were detected in the testes at stages III and VI. Pseudotime analysis showed a development trajectory from stage I through stage VI to III, reflecting the testes at stage I may represent the initial state of testicular development. In fish, testicular development is generally classified into six stages, from I to VI, based on the presence of main and typical spermatogenic cells (Chaves-Pozo et al., 2005; Siqueira-Silva et al., 2013; Schulz et al., 2010; Yan et al., 2022). Typically, the testis at stage I occurs only once in a fish’s lifetime, and is characterized by a large number of spermatogonia (Liu et al., 2021; Schulz et al., 2010; Yang et al., 2018). During spermatogenesis, these SPG undergo meiosis and gradually differentiate into millions of spermatozoa in the spawning season (Schulz et al., 2010; Wang et al., 2023). At postspawning, the testes regress into stage VI, during which diploid spermatogonia predominantly remain as the germ cells capable of initiating the next cycle of spermatogenesis (Chaves-Pozo et al., 2005; Sato et al., 2017; Yu et al., 2024). Previous histological studies have demonstrated that spermatogonia are the main germ cells in both the juvenile and adult regressed testes in many fish species (Sato et al., 2017; Yu et al., 2024). However, our study discovered that the majority of SPG in the adult regressed testes (at stage VI) were Diff SPG rather than Undiff SPG, suggesting differences in stemness between spermatogonia in juvenile and adult regressed testes. This observation aligns with the research in rainbow trout, which demonstrate a higher incorporation efficiency of SPG from prepubertal testes compared to those derived from adult testes using germ cell transplantation method (Sato et al., 2017). This finding offers valuable insight for further research on spermatogonia analysis and germ cell transplantation in teleost. Furthermore, we assessed the expression levels of specific marker genes in the testes at six stages from I to VI, by RT-qPCR analysis. These findings are in accordance with those reported in some other fish species (Lacerda et al., 2019; Lau et al., 2013; Yang et al., 2018), reflecting the spermatogenic cell types and their marker genes are varied significantly depending on the reproductive stages.

### 3. Identification and characteristics of SSC

In vertebrates, from teleost to human, SSCs, a rare subset of spermatogonia, are the origin of spermatogenesis. Identification and characterization of various spermatogonia populations have been reported in several species by scRNA-seq (Guo et al., 2018; Huang et al., 2023; Lau et al., 2020; Li et al., 2022; Qian et al., 2022; Wang et al., 2022; Wei, et al., 2021; Wu et al., 2021). In our research, we provide new insights into the spermatogonia development, most importantly the identification of a novel and early quiescent fish SSCs. In vertebrate, SSCs are a rare subset of spermatogonia, capable of self-renew and also giving rise to progenitors that are poised for differentiation (Kubota et al., 2003; Lacerda et al., 2014; Lord and Nixon, 2020; Schulz et al., 2010). In this study, we found 3 distinct clusters of Undiff SPG by re-clustering the testicular cells from the juvenile testes (at stage I). Somewhat surprisingly, we found that only a small subset of cells in Undiff SPG state 0 showed high expression levels of Undiff SPG markers, such as *pou5f3, cd9*, *klhdc1, ism14b* and *arhgef19*. Moreover, the RNA velocity analysis also revealed that these assumed SSCs displayed long velocity vectors turning into a state that was likely to be stem cells. The findings strongly suggest that the small subset of cells in the Undiff SPG state 0 were likely to represent the SSC population.

To further elucidate the characteristics of SSC, we performed functional enrichment analyses of marker genes of SSCs and Undiff SPG. Surprisingly, we discovered a clear enrichment of genes associated with “mitochondrial matrix” in SSCs, whereas genes in Undiff SPG were enriched in “respiratory chain”. This suggests equivalent changes in metabolism between SSC and Undiff SPG, with SSCs favoring glycolytic metabolism, whereas Undiff SPG preferring oxidative phosphorylation (OXPHOS) for ATP production. ISH and gene expression analysis further revealed that glycolysis-related genes (*myc, aimpa, eno3* and *gaphd*) were highly expressed in SSCs. In teleost, despite the critical importance of SSCs for male fertility, the phenotypic, biochemical or molecular characteristics of SSCs development have remained largely undefined. Recently, the analyses of scRNA-seq databases from mouse and human testes have suggested a metabolic shift from glycolysis to oxidative phosphorylation during SSCs differentiation (Guo et al., 2017; Hermann et al., 2018; Lord and Nixon, 2020). In other multipotent adult stem cells, the correlation between glycolysis and stem cell maintenances has also been demonstrated in mice and human (Hu et al., 2016; Lord and Nixon, 2020). Furthermore, studies in mice have shown that overexpression of *myc* increases the self-renewal capacity of SSCs (Kanatsu-Shinohara et al., 2016). Understanding the biochemical characteristics of SSCs is critical for establishing a long-term expanding culture of SSCs, making them ideal targets for genetic modification and preservation. Although numerous studies have reported cultures of SSCs in vertebrate, there is no optimal system or culture condition for the long-term culture of SSCs in vitro, especially in non-mammalian species. In this study, we have identified the first SSCs population from Undiff SPG in teleost and discovered their glycolytic metabolism, which may provide unique insights for further fish SSCs culture. Additionally, our analysis of DEGs in SSCs identified several potential marker genes for SSCs, including *ly75-like, bzwz, nthl1* and *cd200*. ISH analysis showed that the *ly75-like* and *ox2* were obviously expressed in Undiff SPG at the periphery of the testes. These genes will offer valuable tools for future research on SSCs in teleost.

### 4. Crosstalk between somatic cells and spermatogonia in the juvenile testes

In vertebrates, Sertoli cells and Leydig cells are the most important somatic cells, and are known for their roles in fostering male germ cell differentiation and production of mature sperm. Sertoli cells are the only somatic cells that come into direct contact with the male germ cells, whereas Leydig cells are outside of the seminiferous tubules and are the steroidogenic lineage for steroid hormones, promoting spermatogenesis (DeFalco et al., 2013; Yao and Rodriguez, 2023). In this study, we found that the specific marker genes for Sertoli cells (*gsdf* and *fgf2*) and Leydig cells (*igf2* and *bmp4*) were significantly expressed in a cell cluster from the juvenile testes, suggesting that the Sertoli cells and Leydig cells may be differentiated from the same cell precursor. Although the origin of Sertoli and Leydig cells in fish is not fully understood, the fetal Sertoli cells were well known to induce appearance of Leydig cells in mammals (Imaimatsu et al., 2023; Yao and Rodriguez, 2023). Our findings offer valuable insights for further research into the origin of gonadal somatic cells in fish.

Furthermore, cell communication analysis showed that a robust interaction between Sertoli/Leydig cells and four subsets of Undiff SPG through NOTCH1/2-JAG1 system. The highly conserved NOTCH-JAG system plays an important role in the control of cell fate in invertebrate and vertebrate developmental process (Dirami et al., 2001; Garcia et al., 2017). In mice and human, Notch ligands and Notch receptor are present in spermatogonia and Sertoli cells (Garcia et al., 2017). Further, in vivo experiments demonstrated that Undiff SPG are activating Notch signaling in Sertoli cells through their expression of the Notch receptor Jag1, therefore maintaining spermatogonia stemness by a ligand concentration-dependent process (Parekh et al., 2019). Our results suggested that the Notch-Jag1 system also played an important role in spermatogonia development in teleost. In addition, we found evidence of robust communication in the Sertoli/Leydig cells through four Ligand-receptor pairs, including PTPRK-BMP7, PLXNB2-SEMA4C, NOTCH1-JAG1 and NOTCH2-JAG1.

In summary, our study of single-cell transcriptomes has generated a comprehensive single-cell transcriptional atlas of testicular development in the large yellow croaker. We have identified maker genes for various germ cells and demonstrated stage-specific heterogeneity in fish testicular cells throughout the annual reproductive cycle. Additionally, we discovered a distinct population of SSCs and observed a metabolic shift from glycolysis to oxidative phosphorylation, which deepens our understanding of the biochemical and molecular characteristics of SSCs development in vertebrates (Figure 6B). Our analysis also clarified the interactions between somatic cells and spermatogonia, shedding light on the mechanisms that regulate SSCs development. This work not only enhances our knowledge of germ cell development in vertebrate but also provides valuable insights for the further conservation and in vitro culture of SSCs in teleost.

## Materials and methods

### 1. Samples

All samples of large yellow croakers were obtained from the research station of Zhejiang Maine Fisheries Research Institute (Xishan Island, City of Zhoushan, China). All experiments were approved by the Institutional Animal Care and Use Ethics Committee of Zhejiang Marine Fisheries Research Institute ([2023]21). Throughout the breeding season, we randomly collected 6-8 individuals at each development stage. The males were sampled at the age of 4, 6, 12, 13, 14, 18 and 22 months. In August and November 2021, and January 2022, ∼50 fish of 4-month-old, one fish of 18-month-old and one fish of 22-month-old were maintained for constructing the datasets for scRNA-seq. All samples were anesthetized with MS-222, then the testicular tissues were dissected for the following experiments.

### 2. Testicular cell preparation for single cell RNA sequencing

For each single cell sequencing experiment, tissues were washed with 1× PBS twice. The blood and tunica albuginea of testes were removed on ice under microscopic examination. The testes were cut into small pieces via scissor, then incubated in L-15 medium solution (Gibco Invitrogen) containing 0.25% trypsin, 5% fetal bovine serum, 20mg/ml collagenase H and 0.05% DNase I at 25°C for 3 hours with gentle agitation. The digestions were stopped by adding 10% FBS. The cell suspensions were filtered with strainers with mesh size 150 μm and 50 μm. The cells were pelleted by centrifugation at 200 X g for 5 min, and resuspended in 1× PBS. The cell number were measured using a blood counting chamber with Trypan Blue dying.

### 3. 10× Genomics single cell RNA-seq

The scRNA-seq library preparation and sequencing were carried out at oebiotech Technology Co. ScRNA-seq was performed using 10× Genomics system. Briefly, cells were diluted following manufacture recommendations, mixed with 33.4 μl master mixed buffer, and were loaded into a 10× chromium controller using Chromium single cell 3’ library & single cell 3’ v3 solution. Each sequencing library was prepared following the manufacturer’s instructions. The resulting libraries were then sequenced on an IIIumina hiseq 2500 instrument.

### 4. Processing of single cell RNA-seq data

The Cell Ranger software pipeline (version 3.1.0) provided by 10× Genomics was used to demultiplex cellular barcodes, map reads to the genome and transcriptome using the STAR aligner, and down-sample reads as required to generate normalized aggregate data across samples, producing a matrix of gene counts versus cells. We processed the unique molecular identifier (UMI) count matrix using the R package Seurat (version 3.0). To remove low quality cells and likely multiplet captures, which is a major concern in microdroplet-based experiments, we applied a filter criterion to remove cell data with UMI/gene numbers out of the limit of mean value +/- 2 fold of standard deviations assuming a Guassian distribution of each cells’ UMI/gene numbers. Following visual inspection of the distribution of cells by the fraction of mitochondrial genes expressed, we further discarded low-quality cells where a larger than normal percentage of counts belonged to mitochondrial genes. Library size normalization was performed in Seurat on the filtered matrix to obtain the normalized count.

The top variable genes across single cells were identified. Briefly, the average expression and dispersion were calculated for each gene, and those genes were subsequently placed into several bins based on expression. Principal component analysis (PCA) was performed to reduce the dimensionality on the log transformed gene-barcode matrices of top variable genes. Cells were clustered based on a graph-based clustering approach, and were visualized in 2-dimension using tSNE. Likelihood ratio test that simultaneously test for changes in mean expression and in the percentage of expressed cells was used to identify significantly differentially expressed genes between clusters. Here, we use the R package Single R, a novel computational method for unbiased cell type recognition of scRNA-seq to infer the cell of origin of each of the single cells independently and identify cell types. Differentially expressed genes (DEGs) were identified using the Seurat package. *P value* < 0.05 and |log2foldchange| > 1 was set as the threshold for significantly differential expression. GO enrichment and KEGG pathway enrichment analysis of DEGs were respectively performed using R based on the hypergeometric distribution.

### 5. Histology

After anesthetization, each fish was dissected, then the gonad was fixed in Bouin’s solution for histological analysis and immunohistochemistry, and was fixed in RNAwait (Biosharp, Guangzhou, China) and then stored at −80 °C for quantitative real-time PCR (RT-qPCR). For histological analysis, the fixed gonads were embedded in paraffin, cut into 5-μm-thick sections, and stained with hematoxylin and eosin (HE). The results of HE staining were examined and photographed using light microscopy. More than five views per section were examined for each fish. The germ cell type identification was followed established in previous studies (Yang et al., 2018).

### 6. Immunohistochemistry

The immunofluorescence staining in testicular tissues at different reproductive stages were performed using Vasa antibody (ab209710, Abcam). The testes fixed in Bouin’s solution were embedded in paraffin wax, and then sliced into 5 μm serial sections. Immunohistochemistry was conducted following the detailed protocols of our lab, as described previously (Yu et al., 2024). Fluorescence images were captured with a fluorescence microscope (model BX-51N-34FL, Olympus).

### 7. In *situ* hybridization

The RNA probes for detected genes were synthesized using a DIG RNA Labeling Kit (SP6/T7) (Roche, Germany). ISH was performed on 5 μm 4% paraformaldehyde-fixed paraffin-embedded sections from the testicular tissues. ISH was conducted following the detailed protocols of our lab, as described previously (Yang et al., 2019). The specific primers for detected genes were listed in supplementary Table 1.

### 8. RT-qPCR analysis

The specific primers for detected genes were designed using Primer Premiers of 5.0 and listed in supplementary Table 1. Total RNA was extracted from testes using the RNA extraction kit (Solarbio, China). Genomic DNA removal and cDNA synthesis were performed with a Transcript First-strand cDNA Synthesis Kit (TranStart, China). The RT-qPCR amplifications were performed using 2× SYBY Premix Ex TapTM II (Takara, China) as described previously. Each experiment was performed in triplicates. The relative gene expression levels were analyzed using the 2^−ΔΔCT^ method.

### 9. Statistical analysis

All datas were presented as means ± standard error of the mean (SEM) using GraphPad Prism 9.0 software. Statistical differences were calculated using one-way ANOVA followed by the Kruskal-Waillis multiple comparisons test (*P*< 0.05).

## Data availability

The scRNA-seq data generated in this study have been deposited in GEO under the accession number GSE269124.

---

**Supplementary Figure 1.**
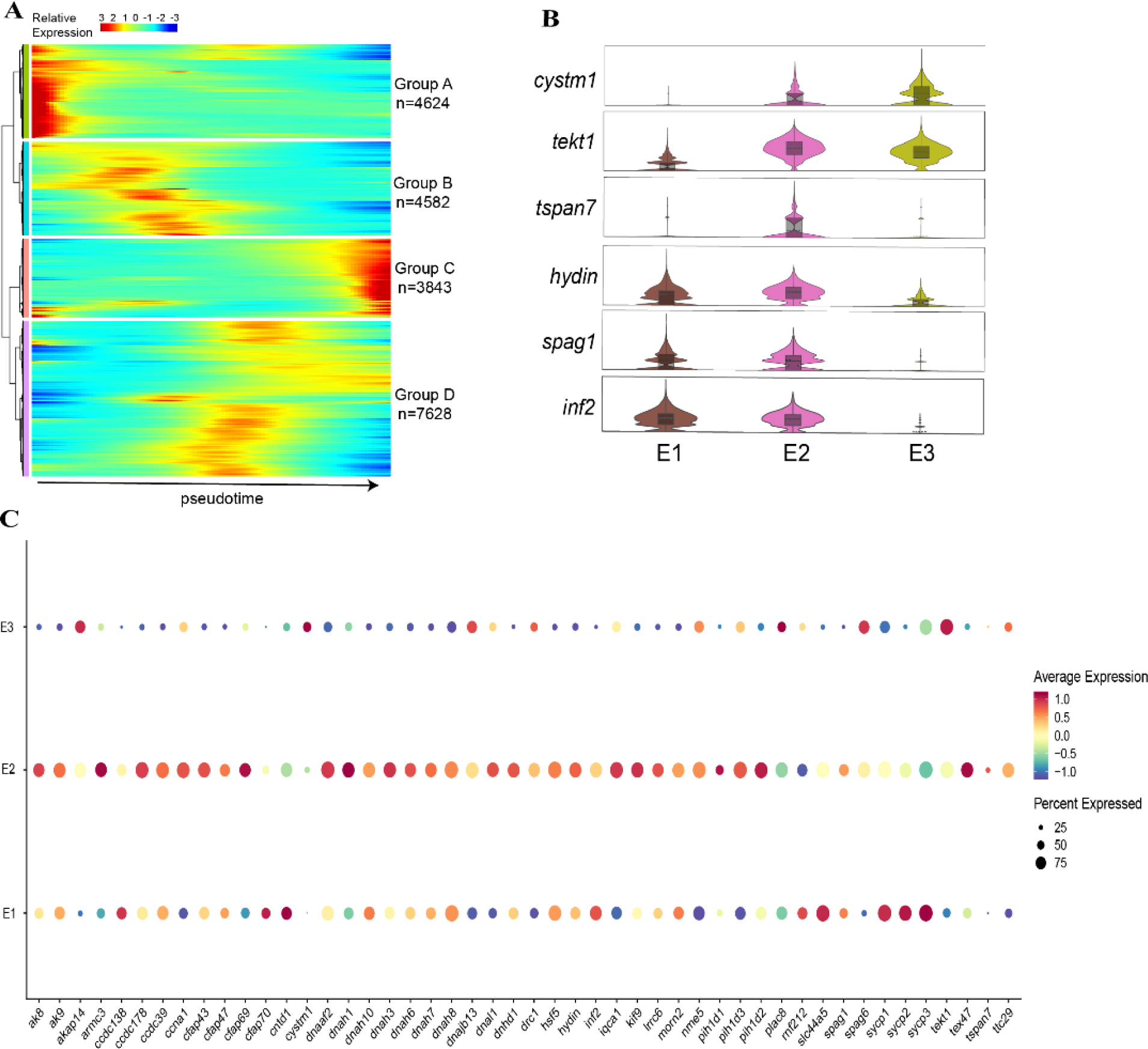
Analysis of spermiogenesis in large yellow croaker. (A) Gene expression heatmap in pseudotime. After a clustering analysis, genes were divided into four groups. (B) Volin plots of selected genes of spermiogenesis in 3 sperm clusters. (C) bubble blots indicate the average expression levels of DEGs in 3 sperm clusters.

**Supplementary Figure 2.**
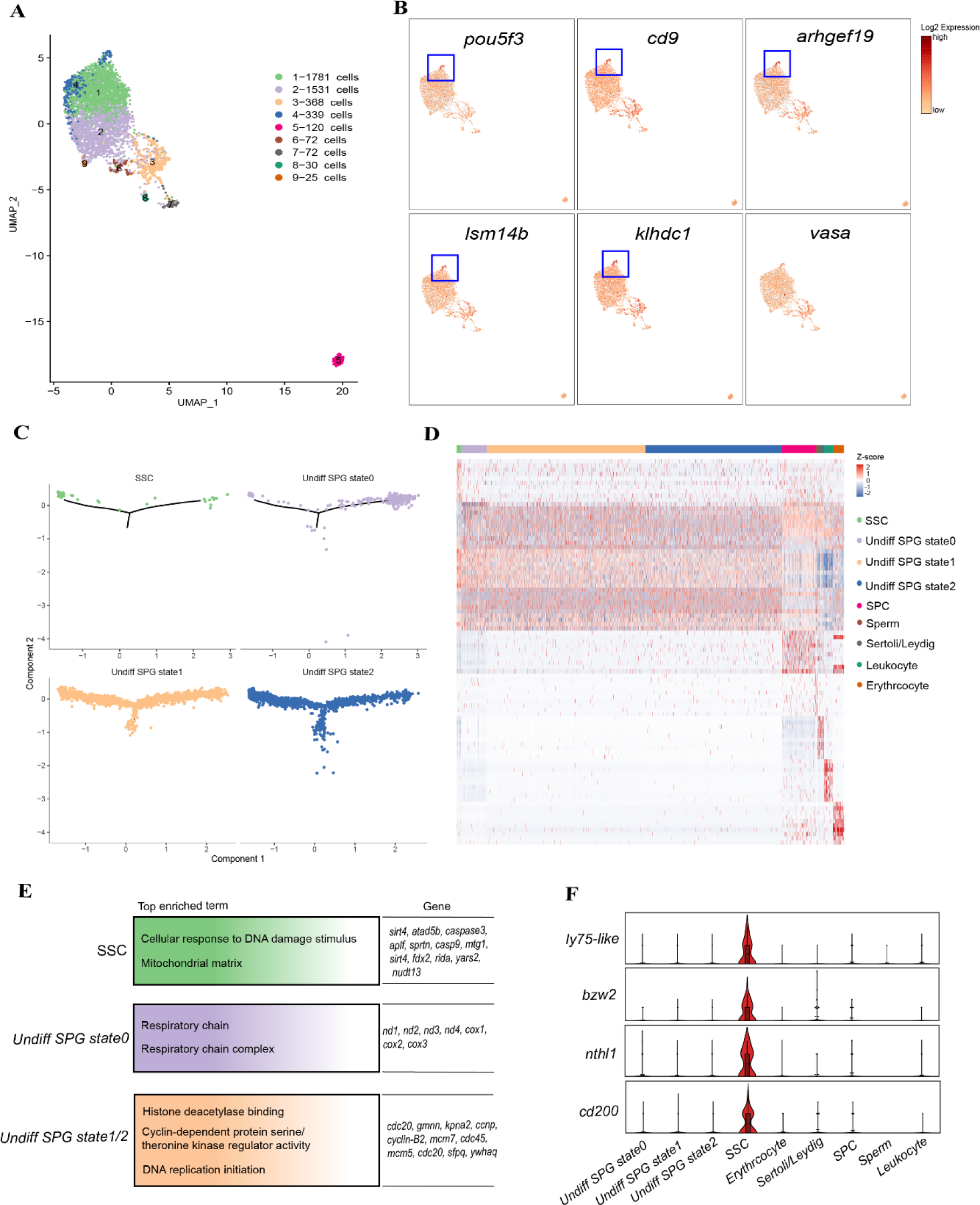
Single-cell transcriptome analysis from juvenile testes (4-months old) of large yellow croaker. (A) A UMAP plot showing 9 cell types from juvenile testes of large yellow croaker. (B) Expression patterns of selected Undiff SPG marker genes on the UMAP plot. Red (or orange) represents a high (or low) expression level. (C) Pseudotime analysis of SSC and Undiff SPG state 0-2. (D) Heatmap showing the top 20 DEGs for 9 cell types. (E) Functional enrichment analysis of DEGs in SSC and Undiff SPG state 0-2. (F) Violin plot showing selected SSC marker genes in 9 cell types. Undiff SPG, undifferentiated spermatogonia; Diff SPG, differentiated spermatogonia; SPC, spermatocyte.

**Supplementary Figure 3.**
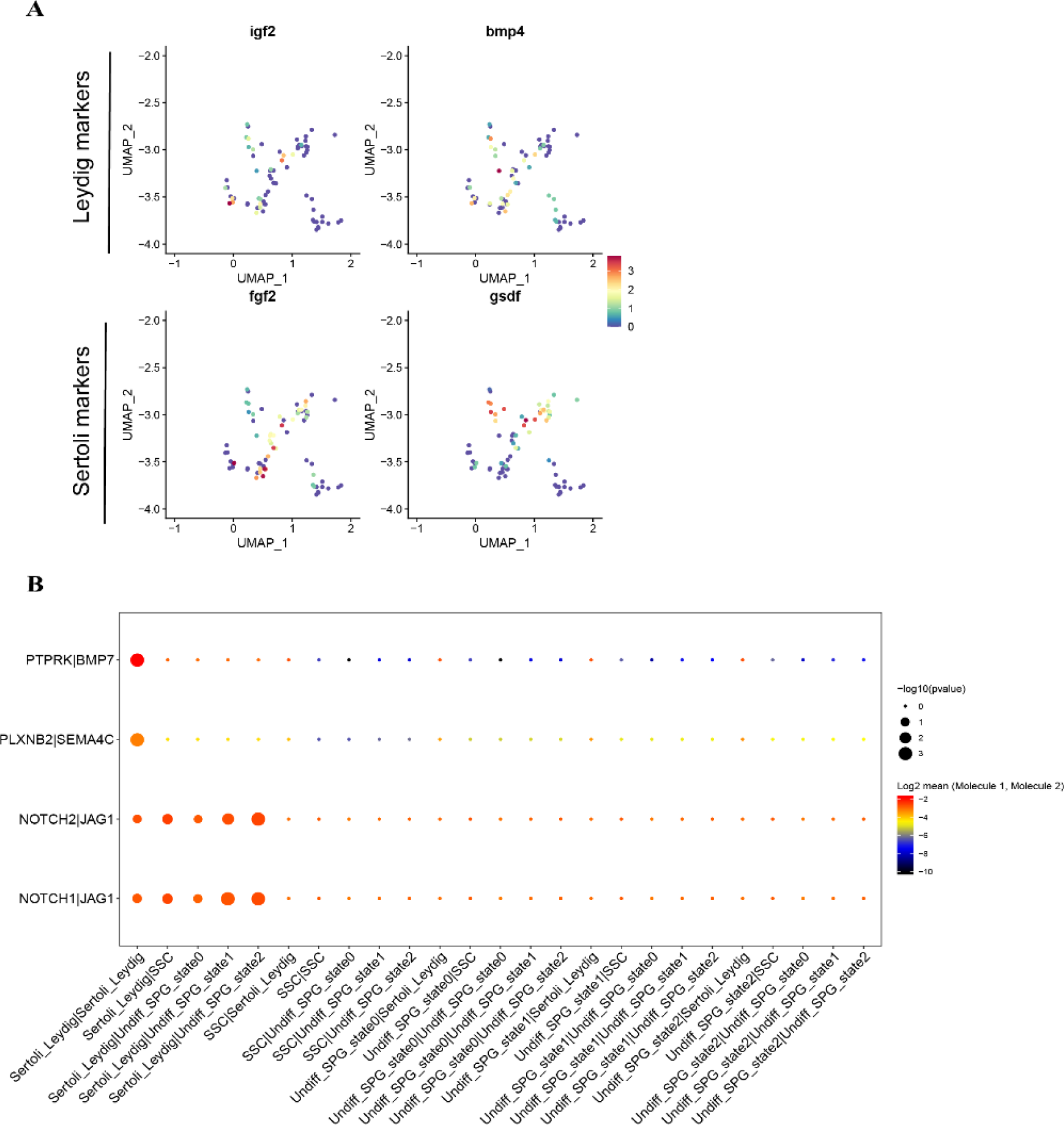
Analysis of crosstalk between somatic cells and spermatogonia of large yellow croaker. (A) A UMAP plot showing selected marker genes in Sertoli/Leydig cells from juvenile testes. (B) bubble plot indicates the significance and average expression levels of ligand-receptor pairs. Undiff SPG, undifferentiated spermatogonia; Diff SPG, differentiated spermatogonia; SPC, spermatocyte.

**Supplementary Table 1.**
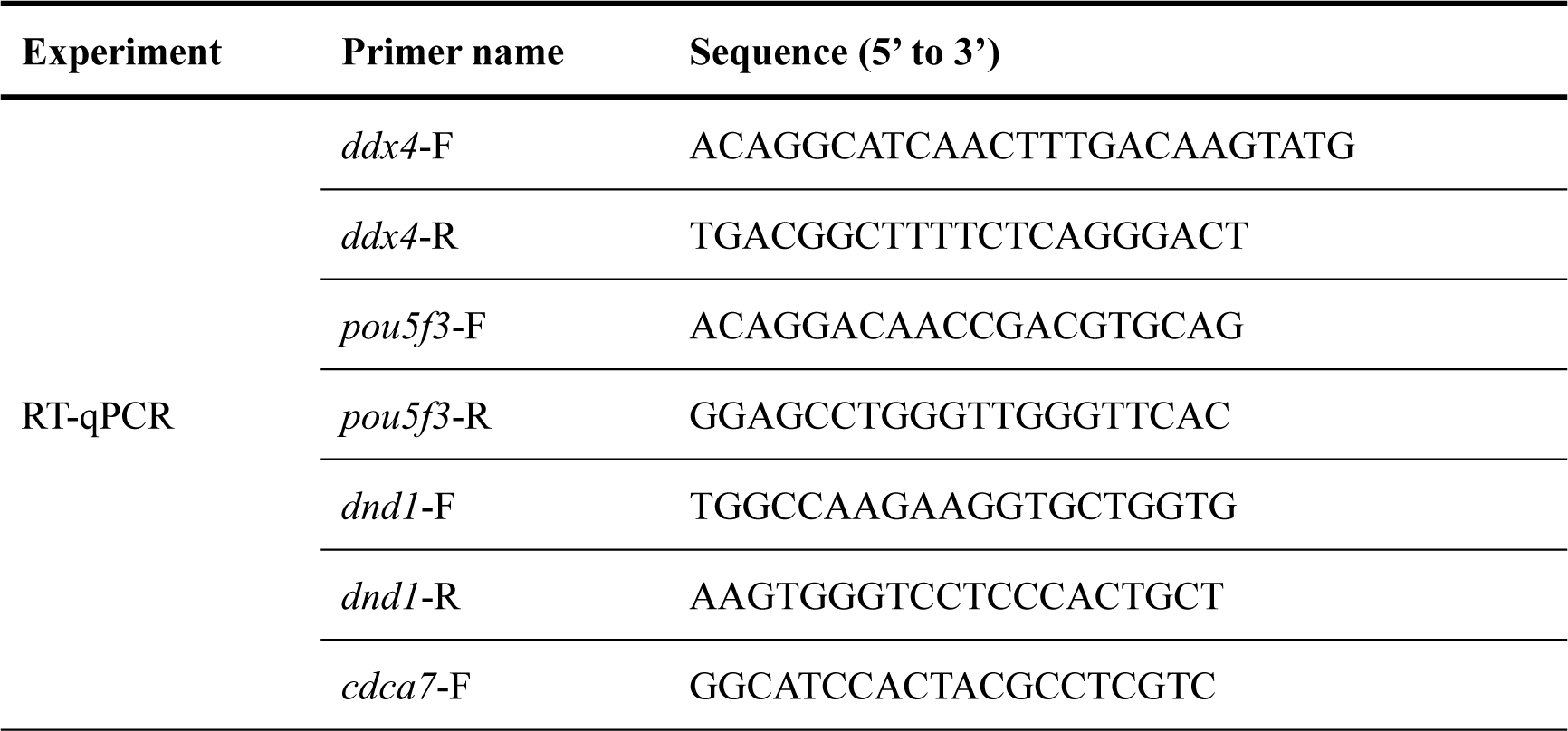
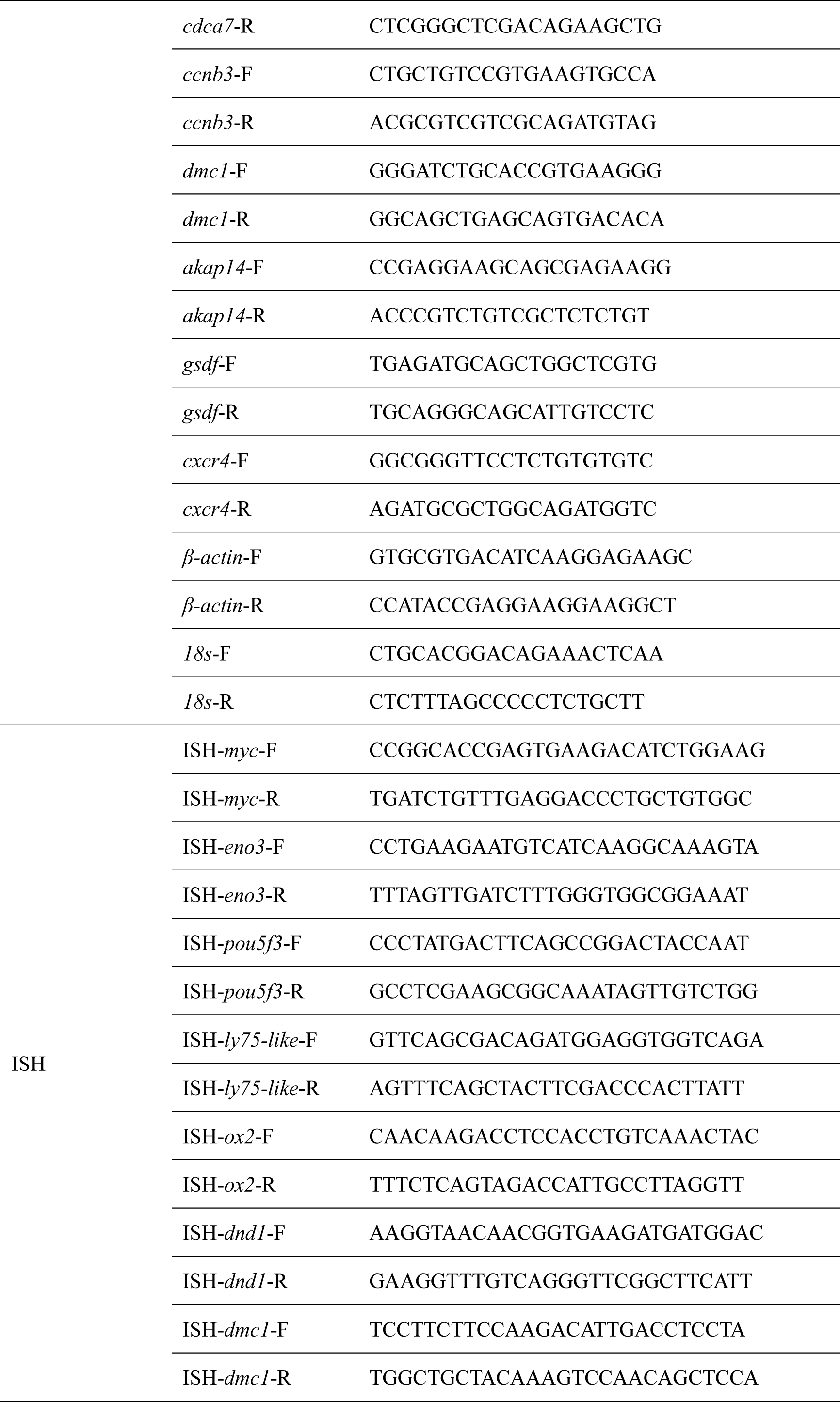
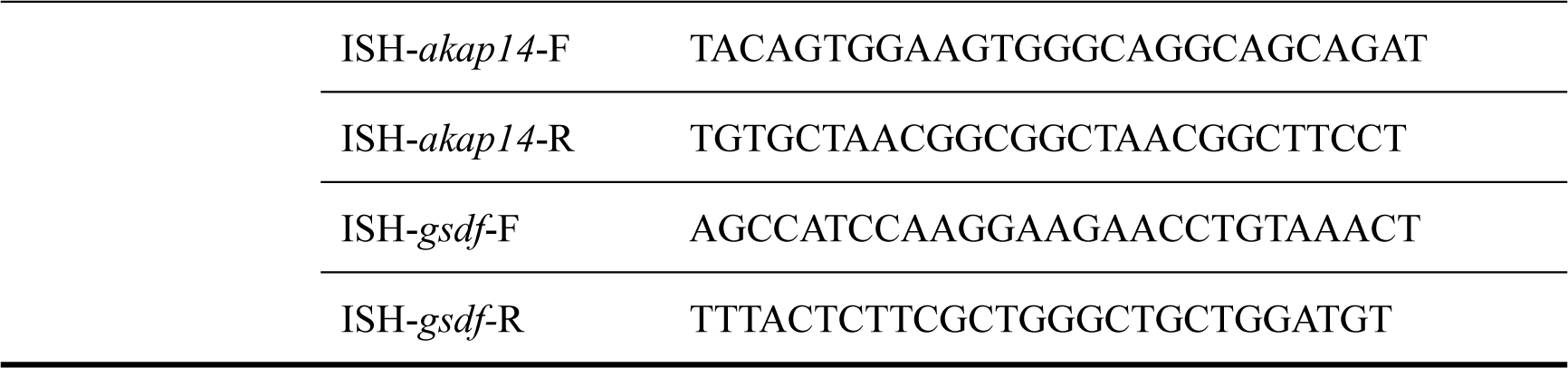
Primers used in this study.

## References

Chaves-Pozo, E., Mulero, V., Meseguer, J., Ayala, A.G. (2005). An overview of cell renewal in the testis throughout the reproductive cycle of a seasonal breeding teleost, the gilthead seabream (*Sparus aurata L*). Biol. Reprod. 72. 593–601.

Chen, S., Su, Y., Hong, W. (2018). Aquaculture of the large yellow croaker. Aquac. China. 297–308.

DeFalco, T., Saraswathula, A., Briot, A., Iruela-Arispe, M.L., Capel, B. (2013). Testosterone levels influence mouse fetal leydig cell progenitors through notch signaling1. Biol. Reprod. 91. 1–12.

Dirami, G., Ravindranath, N., Achi, M.V., Dym, M. (2001). Expression of notch pathway components in spermatogonia and sertoli cells of neonatal mice. J. Androl. 22. 944–952.

Dong, F., Ping, P., Ma, Y., Chen, X.F. (2023). Application of single-cell RNA sequencing on human testicular samples: a comprehensive review. Int. J. Bio. Sci. 19. 2167–2197.

Garcia, T.X., Parekh, P., Gandhi, P., Sinha, K., Hofmann, M.-C. (2017). The notch ligand jag1 regulates gdnf expression in sertoli cells. Stem Cells Dev. 26. 585–598.

Guo, J., Grow, E.J., Mlcochova, H., Maher, G.J., Lindskog, C., Nie, X., Guo, Y., Takei, Y., Yun, J., Cai, L., Kim, R., Carrell, D.T., Goriely, A., Hotaling, J.M., Cairns, B.R. (2018). The adult human testis transcriptional cell atlas. Cell. Res. 28. 1141–1157.

Guo, J., Grow, E.J., Yi, C., Mlcochova, H., Maher, G.J., Lindskog, C., Murphy, P.J., Wike, C.L., Carrell, D.T., Goriely, A., Hotaling, J.M., Cairns, B.R. (2017). Chromatin and single-cell RNA-seq profiling reveal dynamic signaling and metabolic transitions during human spermatogonial stem cell development. Cell Stem Cell. 21. 533–546

Hermann, B.P., Cheng, K., Singh, A., Roa-De La Cruz, L., Mutoji, K.N., Chen, I.C., Gildersleeve, H., Lehle, J.D., Mayo, M., Westernströer, B., Law, N.C., Oatley, M.J., Velte, E.K., Niedenberger, B.A., Fritze, D., Silber, S., Geyer, C.B., Oatley, J.M., McCarrey, J.R. (2018). The mammalian spermatogenesis single-cell transcriptome, from spermatogonial stem cells to spermatids. Cell. Rep. 25. 1650–1667.

Hernández-Franyutti, A., Uribe, M.C. (2012). Seasonal spermatogenic cycle and morphology of germ cells in the viviparous lizard *Mabuya brachypoda* (Squamata, Scincidae). J. Morphol. 273. 1199–1213.

Hu, C., Fan, L., Cen, P., Chen, E., Jiang, Z., Li, L. (2016). Energy metabolism plays a critical role in stem cell maintenance and differentiation, Int. J. Mol. Sci.17(2). 253.

Huang, L., Zhang, J., Zhang, P., Huang, X., Yang, W., Liu, R., Sun, Q., Lu, Y., Zhang, M., Fu, Q. (2023). Single-cell RNA sequencing uncovers dynamic roadmap and cell-cell communication during buffalo spermatogenesis. iScience. 26. 105733.

Imaimatsu, K., Hiramatsu, R., Tomita, A., Itabashi, H., Kanai, Y. (2023). Partial male-to-female reprogramming of mouse fetal testis by Sertoli cell ablation. Development. 150(14). 201660.

Itman, C., Loveland, K.L. (2013). Smads and cell fate: Distinct roles in specification, development, and tumorigenesis in the testis. IUBMB Life. 65. 85–97.

Kanatsu-Shinohara, M., Tanaka, T., Ogonuki, N., Ogura, A., Morimoto, H., Cheng, P.F., Eisenman, R.N., Trumpp, A., Shinohara, T. (2016). Myc/Mycn-mediated glycolysis enhances mouse spermatogonial stem cell self-renewal. Genes & Dev. 30. 2637–2648.

Kanatsu-Shinohara, M., Toyokuni, S., Shinohara, T. (2004). CD9 is a surface marker on mouse and rat male germline stem cells. Biol. Reprod. 70(1). 70–75.

Kubota, H., Avarbock, M.R., Brinster, R.L., 2003. Spermatogonial stem cells share some, but not all, phenotypic and functional characteristics with other stem cells. PNAS. 100. 6487–6492.

Kubota, H., Brinster, R.L. (2018). Spermatogonial stem cells. Biol. Reprod. 99. 52–74.

Lacerda, S.M.D.S.N, Costa, G.M.J, de França, L.R. (2014). Biology and identity of fish spermatogonial stem cell. Gen. Comp. Endocrinol. 207. 56–65.

Lacerda, S.M.S.N., Martinez, E.R.M., Mura, I.L.D.D., Doretto, L.B., Costa, G.M.J., Silva, M.A., Digmayer, M., Nóbrega, R.H., França, L.R. (2019). Duration of spermatogenesis and identification of spermatogonial stem cell markers in a Neotropical catfish, Jundiá (*Rhamdia quelen*). Gen. Comp. Endocrinol. 273. 249–259.

Lau, E.L., Lee, M.F., Chang, C.F. (2013). Conserved sex-specific timing of meiotic initiation during sex differentiation in the protandrous black porgy *Acanthopagrus schlegelii*. Biol. Reprod 88(6), 1–13.

Lau, X., Munusamy, P., Ng, M.J., Sangrithi, M. (2020). Single-cell RNA sequencing of the *Cynomolgus* Macaque testis reveals conserved transcriptional profiles during mammalian spermatogenesis. Dev. Cell. 54. 548–566.

Levkova, M., Radanova, M., Angelova, L. (2022). Potential role of dynein-related genes in the etiology of male infertility: A systematic review and a meta-analysis. Andrology. 10. 1484–1499.

Liu, Y., Liu, Q., Xu, S., Wang, Y., Feng, C., Zhao, C., Song, Z., Li, J. (2021). A deep insight of spermatogenesis and hormone levels of aqua-cultured turbot (*Scophthalmus maximus*). Front. Mar. Sci. 7. 592880.

Lord, T., Nixon, B. (2020). Metabolic changes accompanying spermatogonial stem cell differentiation. Dev. Cell. 52. 399–411.

Niimi, Y., Imai, A., Nishimura, H., Yui, K., Kikuchi, A., Koike, H., Saga, Y., Suzuki, A. (2019). Essential role of mouse Dead end1 in the maintenance of spermatogonia. Dev. Biol. 445. 103-112.

Oatley, J.M., Brinster, R.L. (2006). Spermatogonial Stem Cells. Methods Enzymol. 419. 259–282.

Parekh, P.A., Garcia, T.X., Waheeb, R., Jain, V., Gandhi, P., Meistrich, M.L., Shetty, G., Hofmann, M.C. (2019). Undifferentiated spermatogonia regulate *Cyp26b1* expression through NOTCH signaling and drive germ cell differentiation. FASEB. J. 33. 8423–8435.

Persio, S.D, Neuhaus, N. (2023). Human spermatogonial stem cells and their niche in male (in) fertility: novel concepts from single-cell RNA-sequencing. Hum. Reprod. 38(1). 1–13.

Priyanka, P.P., Yenugu, S. (2021). Coiled-coil domain-containing (CCDC) proteins: Functional roles in general and male reproductive physiology. Reprod. Sci. 28. 2725–2734.

Qian, P., Kang, J., Liu, D., Xie, G. (2022). Single cell transcriptome sequencing of zebrafish testis revealed novel spermatogenesis marker genes and stronger Leydig-germ cell paracrine interactions. 13. 851719

Sato, M., Hayashi, M., Yoshizaki, G. (2017). Stem cell activity of type A spermatogonia is seasonally regulated in rainbow trout. Biol. Reprod. 96. 1303–1316.

Schulz, R.W., de Franca, L.R., Lareyre, J.J., LeGac, F., Chiarini-Garcia, H., Nobrega, R.H., Miura, T. (2010). Spermatogenesis in fish. Gen. Comp. Endocrinol. 165. 390–411.

Seki, S., Kusano, K., Lee, S., Iwasaki, Y., Yagisawa, M., Ishida, M., Hiratsuka, T., Sasado, T., Naruse, K., Yoshizaki, G. (2017). Production of the medaka derived from vitrified whole testes by germ cell transplantation. Sci. Rep. 7. 43185–43185.

Siqueira-Silva, D.H.D, Vicentini, C. A., Ninhaus-Silveira, A., Veríssimo-Silveira, R. (2013). Reproductive cycle of the Neotropical cichlid yellow peacock bass *Cichla kelberi*: A novel pattern of testicular development. Neotrop. Ichthyol. 11(3). 587–596.

Tang, S., Wang, X., Li, W., Yang, X., Li, Z., Liu, W., Li, C., Zhu, Z., Wang, L., Wang, J., Zhang, L., Sun, X., Zhi, E., Wang, H., Li, H., Jin, L., Luo, Y., Wang, J., Yang, S., Zhang, F. (2017). Biallelic mutations in CFAP43 and CFAP44 cause male infertility with multiple morphological abnormalities of the sperm flagella. Am. J. Hum. Genet. 100. 854–864.

von Kopylow, K., Spiess, A.-N. (2017). Human spermatogonial markers. Stem Cell Res. 25. 300–309.

Wang, H. Y., Liu, X., Chen, J.-Y., Huang, Y., Lu, Y., Tan, F., Liu, Q., Yang, M., Li, S., Zhang, X., Qin, Y., Ma, W., Yang, Y., Meng, L., Liu, K., Wang, Q., Fan, G., Nóbrega, R.H., Liu, S., Piferrer, F., Shao, C. (2022). Single-cell-resolution transcriptome map revealed novel genes involved in testicular germ cell progression and somatic cells specification in Chinese tongue sole with sex reversal. Sci China Life Sci. 66. 1151–1169.

Wang, X., Liu, Q., Li, J., Zhou, L., Wang, T., Zhao, N. (2023). Dynamic cellular and molecular characteristics of spermatogenesis in the viviparous marine teleost *Sebastes schlegelii*. Biol. Reprod. 108. 338–352.

Wei, Y.X., Tong, L.T., Min, D.X., Shen, Q.Y., Zhang, M.F., Wei, Y.D., Yang, D.H., Xu, W.J., Chen, W.B., Bai, C.L., Li, X.L., Li, G.P., Li, N., Peng, S., Liao, M.Z., Hua, J.L. (2021). Single-cell RNA sequencing reveals atlas of dairy goat testis cells. Zool. Res. 42. 401–405.

Wu, C., Zhang, D., Kan, M., Lv, Z., Zhu, A., Su, Y., Zhou, D., Zhang, J., Zhang, Z., Xu, M., Jiang, L., Guo, B., Wang, T., Chi, C., Mao, Y., Zhou, J., Yu, X., Wang, H., Weng, X., Jin, J.G., Ye, J., He, L., Liu, Y. (2014). The draft genome of the large yellow croaker reveals well-developed innate immunity. Nat. Commun. 5. 5227.

Wu, X., Yang, Y., Zhong, C., Wang, T., Deng, Y., Huang, H., Lin, H., Meng, Z., Liu, X. (2021). Single-cell atlas of adult testis in protogynous hermaphroditic orange-spotted grouper, *Epinephelus coioides*, Int. J. Mol. Sci. 22(22). 12607.

Xu, D., Yoshino, T., Konishi, J., Yoshikawa, H., Ino, Y., Yazawa, R., Dos Santos Nassif Lacerda, S.M., de França, L.R., Takeuchi, Y. (2019). Germ cell-less hybrid fish: ideal recipient for spermatogonial transplantation for the rapid production of donor-derived sperm. Biol. Reprod. 101. 492–500.

Yan, L.T., Jiang, Y., Xu, Q., Ding, G.-m., Chen, X.Y, Liu, M. (2022). Reproductive dynamics of the Large Yellow Croaker *Larimichthys crocea* (Sciaenidae), a commercially important fishery species in China. Fron. Mar. Sci. 9. 868580.

Yang, Y., Liu, Q., Ma, D., Xiao, Y., Xu, S., Wang, X., Song, Z., You, F., Li, J. (2018). Spermatogonial stem cells differentiation and testicular lobules formation in a seasonal breeding teleost: The evidence from the heat-induced masculinization of genetically female Japanese flounder (*Paralichthys olivaceus*). Theriogenology. 120. 68–78.

Yang, Y., Liu, Q., Xiao, Y., Xu, S., Wang, X., Yang, J., Song, Z., You, F., Li, J. (2019). High temperature increases the *gsdf* expression in masculinization of genetically female Japanese flounder (*Paralichthys olivaceus*). Gen. Comp. Endocrinol. 274. 17–25.

Yao, H.H.C., Rodriguez, K.F. (2023). From Enrico Sertoli to freemartinism: the many phases of the master testis-determining cell. Biol. Reprod. 108. 866–870.

Yokota, Y. (2001). Id and development. Oncogene. 20. 8290–8298.

Yoshida, S. (2016). From cyst to tubule: innovations in vertebrate spermatogenesis. WIREs Dev. Biol. 5. 119–131.

Yu, L., Yang, Y., Yu, Y., Li, H., Chen, R., Miao, L., Xu, D. (2024). Gametogenesis and vasa expression are seasonally regulated in yellow drum (*Nibea albiflora*). Aquacult. Rep. 35. 101970.

Yu, Y., Yang, Y., Ye, H., Lu, L., Li, H., Xu, Z., Li, W., Yin, X., Xu, D. (2023). Identification of germ cells in large yellow croaker (*Larimichthys crocea*) and yellow drum (*Nibea albiflora*) using RT-PCR and in situ hybridization analyses. Gene. 863. 147280.

Zhou, L., Wang, X., Liu, Q., Yang, J., Xu, S., Wu, Z., Wang, Y., You, F., Song, Z., Li, J. (2021). Successful spermatogonial stem cells transplantation within *Pleuronectiformes*: first breakthrough at inter-family level in marine fish. Int. J. Biol. Sci. 17. 4426–4441.

